# Cross-kingdom degradomics identifies natural plant small RNAs with disease protecting activity

**DOI:** 10.1101/2025.10.06.680647

**Authors:** Bernhard T. Werner, Sabrine Nasfi, Tara Procida-Kowalski, Rebekka Schmidt, Jochen Wilhelm, Patrick Schäfer

## Abstract

RNA interference (RNAi) is a highly specific process regulating genes transcriptionally or post-transcriptionally via silencing. small regulatory RNAs (sRNAs) provide the specificity of post-transcriptional silencing by binding to complementary mRNAs leading to their degradation or translational repression. sRNAs are promising non-chemical control agents against plant pests and diseases. Their application requires, however, a deep understanding of the determinants of functional specificity and efficiency. In nature, interacting plants and pathogens follow the manipulative strategy of cross-kingdom RNAi (ckRNAi) based on the targeted transfer of sRNAs. In this study, we employed a genome-wide approach to explore the diversity of natural ck-sRNAs exchanged in the pathogenic interaction of *Brachypodium distachyon* with the fungal pathogen *Fusarium graminearum*. By deep-sequencing of the cross-kingdom sRNAome, transcriptome and degradome 258 ckRNAi-mediating sRNAs were discovered. In a simulation-based approach, specific sequence characteristics allowed categorizing sRNAs into functionality classes. Subsequent genome annotation analyses revealed the organization of plant ck-sRNAs in long non-coding RNAs. Moreover, comparative sequence analyses revealed an evolution of ck-sRNAs towards invariant mRNA target regions. Functional analyses with sRNA candidates confirmed an antipathogenic activity at exceptionally low application doses. It indicates natural sRNA as untapped resource and potential blueprint for the development of highly specific and effective plant-derived bioprotectants.

## Introduction

RNA interference (RNAi) regulates a wide range of fundamental biological processes in plants, fungi and most other eukaryotes such as growth, development, stress adaptation, and immunity (Ketting, 2011). A well characterized RNAi mechanism is post-transcriptional gene silencing (PTGS), which relies on small regulatory RNAs (sRNAs) that silence complementary mRNAs (Hannon, 2002). Argonaute (AGO) proteins are key mediators of PTGS which load sRNAs and become part of a larger RNA-induced silencing complex (RISC). The sRNAs guide the RISC to respective target sites of complementary mRNAs to either cleave mRNAs or repress the translation of the gene product, followed by mRNA decay (Fang & Qi, 2016; Torres-Martínez & Ruiz-Vázquez, 2017).

PTGS has been studied in many species since its mechanistic description in the 1990s (Fire et al. 1998) and has been utilized as an analytic tool even before. By introducing double-stranded RNAs (dsRNAs) into eukaryotic cells, genes can be specifically silenced to study their function in cellular processes or to validate their properties as therapeutic agents (Dykxhoorn & Lieberman, 2005). Furthermore, the role of RNAi in crop protection has been analysed since early 2000 (Borovsky, 2005). The high specificity of sRNAs allows highly specific targeting of organisms. Together with their short design-to-synthesis cycle sRNAs are advantageous in many aspects over chemical crop protection agents. This becomes even more relevant as the repertoire and modes of action of pesticides become limited and their repeated application fosters pesticide resistance in pathogens, further reducing their effectiveness (Koch & Kogel, 2014).

In 2010, Nowara et al. postulated the transfer of plant double-stranded (ds) and antisense RNA to target powdery mildew transcripts and weaken fungal development. Later, plant pathogenic *Botrytis cinerea* was found to translocate sRNAs into its host plant to suppress plant immunity (Weiberg et al., 2013). This process of sRNA exchange was defined as cross-kingdom RNAi (ckRNAi) and observed to be common in pathogenic and mutualistic interactions of all kingdoms (Zeng et al., 2019; Qiao et al., 2023). As reported for *A. thaliana,* plants might further utilize secondary trans-acting small interfering RNAs (tasiRNAs) to silence genes in the pathogenic oomycete *Phytophthora capsici* (Hou et al., 2019). First quantitative screens revealed the translocation of 28 cotton miRNAs into the pathogenic fungus *Verticillium dahliae* (Zhang et al., 2016). Two of these cotton miRNAs downregulated predicted fungal targets and fungal knock-out mutants were less virulent. The development of parallel analysis of RNA ends (PARE) allowed genome-wide sRNA-mediated cleavage site detection and, hence, provided direct evidence of RNAi (German et al., 2009). Based on PARE, 8 wheat sRNAs were identified to target and silence transcripts of the rust pathogen *Puccinia striiformis* (Mueth & Hulbert, 2022). All sRNAs identified in these different screens represented anti-pathogenic agents and have highlighted the potential of sRNAs for future application in the field.

Currently, there are two application strategies for RNAi in crop protection: host-induced gene silencing (HIGS) and spray-induced gene silencing (SIGS) (Koch et al., 2019). HIGS is a transgenic approach where dsRNAs or sRNAs are expressed *in planta*. The strategy was successfully applied to a diversity of pathogens and pests (Koch & Wassenegger, 2021). In SIGS, dsRNAs are directly applied to plants, allowing dsRNA screening without generating genetically modified organisms (Koch et al., 2019). For HIGS and SIGS approaches, the application of custom-designed dsRNAs against predicted targets in pathogens is the common strategy, while naturally evolved plant sRNAs have neither been systematically identified nor applied. Especially, naturally evolved plant sRNAs with anti-fungal activities could serve as a blueprint for SIGS-based plant protection strategies (Cai et al., 2018a).

In this study, we combined mRNA-seq, sRNA-seq, and Degradome-seq to identify natural anti-fungal sRNAs produced in the grass model species *Brachypodium distachyon* that target and degrade transcripts of the notorious pathogen *Fusarium graminearum*. Establishing a highly reproducible ckRNAi analysis pipeline allowed us to identify 197 plant cross-kingdom (ck)-sRNAs and to further categorize them based on specific sequence characteristics. In a second step, we developed an independent platform to accurately quantify the slicing activity of identified ck-sRNA candidates prior to functional SIGS validation assays. Our study indicated, for the first time, the highly effective plant protection capacity of natural ck-sRNAs against *F. graminearum*, their genomic organization in long-noncoding RNAs (lncRNAs), and that their high specificity is based on the co-evolution of ck-RNAs against conserved mRNA target sites. The analyses highlight plant-derived ck-sRNAs as a yet untapped resources for the development of highly potent and specific anti-pathogenic agents for future field applications.

## Results

### High quality sequencing data revealed differentially regulated mRNAs and small RNAs

To study ckRNAi events during the pathogenic interaction of *B. distachyon 21-3* (*Bd21-3*) and *F. graminearum-PH1* (*Fg-PH1*) we inoculated or mock-treated *Bd21-3* seedlings. 3 days post inoculation (dpi) plants displayed evenly distributed necrotic lesions on all leaves and the lower shoots, while control plants remained healthy (Fig. 1A & B). To assess the extent of RNAi the third leaf of infected and control plants was harvested from three independent biological experiments for RNA extraction. To guarantee comparability of data sets, the same RNA samples were used for the generation of mRNA-seq, sRNA-seq and Degradome-seq libraries. A total of 626 million paired end mRNA reads (2 x 51 bp) with a mean quality > Q34 were generated (Tab. S1). mRNA reads were aligned against the *Bd21-3* (BdistachyonBd21_3_537_v1.2) and *Fg-PH1* (GCF_000240135.3_ASM24013v3) reference genomes. Per infected sample, 3.74%, 3.50%, and 3.78%, respectively, of all reads mapped concordantly to *Fg-PH1*, while only 0.04%, 0.03%, and 0.04%, respectively, of reads from the mock-treated samples mapped to *Fg-PH1*, showing an even infection between samples and insignificant miss-mapping of *Bd21-3* mRNA (Tab. S2).

**Figure 1:**
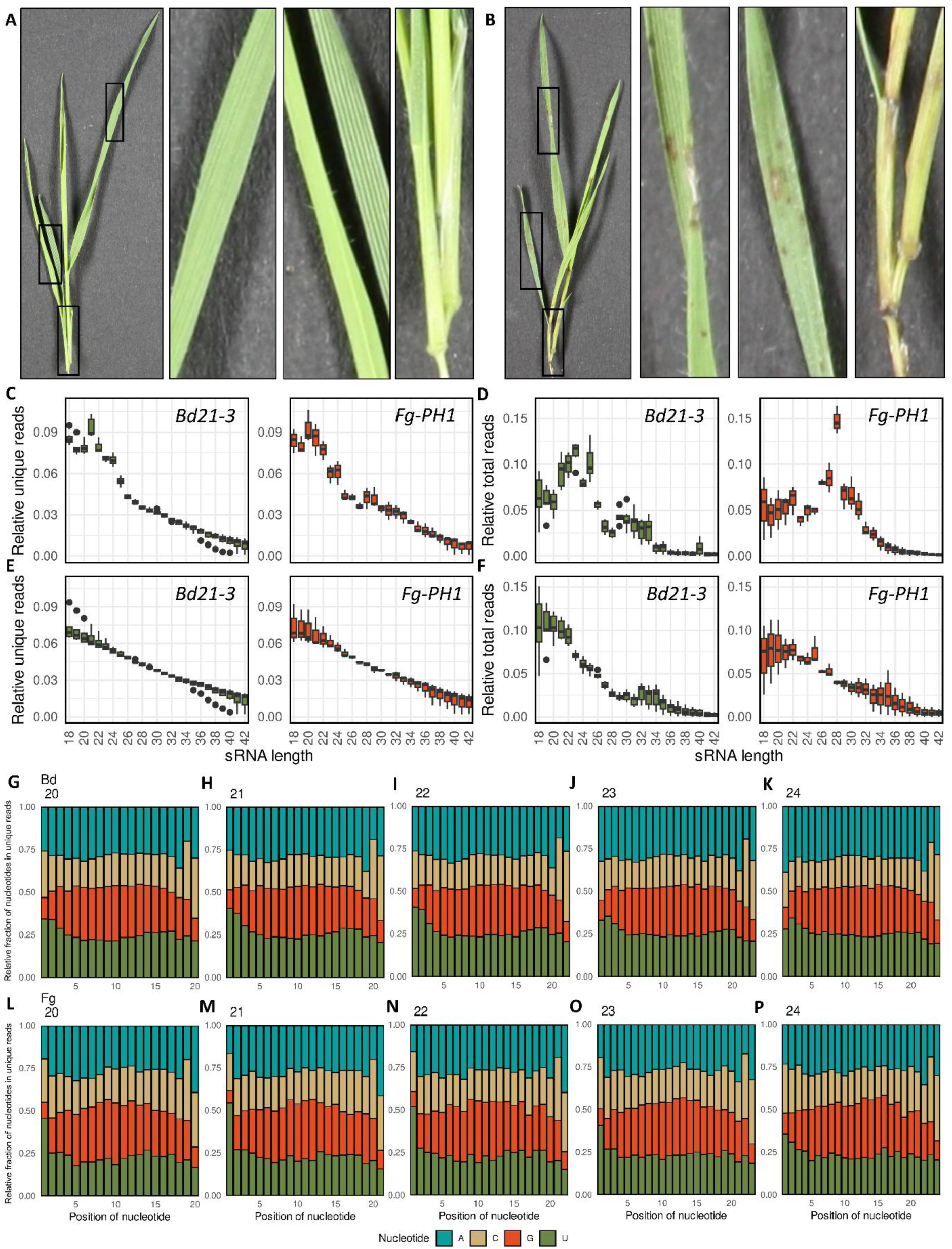
*B. distachyon 21-3* (*Bd21-3*) and *F. graminearum PH1* (*Fg-PH1*) sRNAs from infected and control plants. (A-B) Disease symptoms of *Fg-PH1*-inoculated *Bd21-3* seedlings. (A) Mock-treated seedling leaves at 3 dpt. (B) Symptoms of *Fg-PH1-*inoculated seedling leaves at 3 dpi. (C-F) Relative length distribution of sRNAs. The relative length distribution of (C) non-ribosomal unique reads, (D) non-ribosomal total reads, (E) ribosomal unique reads, (F) ribosomal total reads. sRNA-reads are shown for each sRNA sequencing sample separated by organism of origin. (G-P) Relative fraction of nucleotides per position in unique sRNAs. The nucleotide distributions of unique sRNA reads are shown per length classes 20-24 nt for *Bd21-3* (G-K) and *Fg-PH1* (L-P).

mRNA-seq data analyses revealed 416 significantly up- and 412 downregulated genes (Tab. S3), confirming transcriptional changes in *Bd21-3* in response to the pathogen. Interestingly, a gene ontology enrichment analysis identified upregulated genes involved in oxidation-reduction processes, tryptophan and other amino acid biosynthetic processes, as well as chitinase activity. These were mainly found in the extracellular or apoplastic space or at the ribosomes (Tab. S4) and included *BdiBd21-3.3G0213900*, *BdiBd21-3.4G0163400* (a *short-chain dehydrogenase reductase 3 family protein*), and *BdiBd21-3.2G0520800* (a *histone-lysine N-methyltransferase trithorax-like protein*) as most highly induced genes (> 200 fold-change [FC] expression) (Tab. S3). GO enrichment further showed a general downregulation of photosynthesis, abscisic acid, and cellulose biosynthetic processes (Tab. S5) with *BdiBd21-3.5G0336601* (16 FC), *BdiBd21-3.3G0684100* (a *dehydration-responsive element-binding protein 1D*, 15.8 FC), and *BdiBd21-3.2G0477900* (a *homologue of Arabidopsis thaliana Aluminium sensitive 3*, 15.8 FC) as most strongly suppressed genes.

sRNA reads were aligned to the transcriptomes and genomes of *Bd21-3* and *Fg-PH1*. Unique sRNA reads of 18–42 nucleotides (nt) in length were assigned to each organism if they were represented by at least 20 reads per million (rpm) in any dataset and mapped perfectly and uniquely to only one of the two organisms. Based on our stringent quality setting we identified 2609 *Bd*-sRNAs induced and 202 suppressed in *Fg-PH1*-colonised plants.

### *Fg-PH1* and *Bd21-3* small RNAs displayed distinct sequence characteristics

The length of total (entirety) and unique (distinct) non-ribosomal reads from *Bd21-3* and *Fg-PH1* showed distinct patterns. For *Bd21-3*, a high abundance for unique sRNAs with 21, 22 and 24 nucleotides (nt) in length was detected (after normalization with *Bd21-3* ribosomal unique reads; Fig. 1C & E); a pattern well-known from other plants, representing classes of sRNAs typically involved in PTGS-associated slicing. For *Fg-PH1*, overrepresented length classes reached from 20–24 nt (normalized to *Fg-PH1* ribosomal unique reads; Fig. 1C & E), with the highest abundance for 20 nt sRNAs. Interestingly, *Bd21-3* further produced a high abundance of 25 and *Fg-PH1* of 28 nt sRNAs, respectively (Fig. 1D). Over 30% of the 28 nt reads from *Fg-PH1* originated from two tRNA fragments of the tRF-5c class derived from tRNA-Glu (18.15%, 13.04%, 19.08%) and tRNA-Asp (13.84%, 17.06%, 17.20%). Read mapping to ribosomal sequences from both species showed, as expected, no distinct features (Fig. 1E & F).

In addition to these length distributions, we observed differences in sequence patterns. sRNAs typically associated with Argonaute proteins for gene silencing have a length of 20–25 nt (Borges & Martienssen, 2015). In Arabidopsis, the class of 21 nt was found to have a pronounced preference for a 5’-terminal uracil which facilitates specific Argonaute loading (Mi et al., 2008). This 5’-uracil bias was seen for most size classes in both organisms (Fig. 1G–P) and was most distinct for 21 and 22 nt *Fg-PH1* sRNAs (Fig. 1M & N) with >50% of reads having a 5’-terminal uracil. For *Bd21-3*, this bias was less obvious, though still overrepresented, especially for 21 and 22 nt sRNAs (Fig. 1H & I).

### Improved degradome analytic pipeline for high precision identification of active sRNAs

PTGS-based slicing leaves two mRNA halves. For precise sequencing of the sliced 3’-mRNA fragments, we improved a PARE-seq protocol (Li et al. 2019) to produce a Degradome-seq data set with higher sensitivity for slicing events. To identify ckRNAi events a bioinformatics workflow was developed to recognize mRNA slice sites and to identify responsible sRNAs, under consideration of cross-kingdom RNA transfer.

Sequenced sRNAs that passed our quality filtering were aligned to *Bd21-3* and *Fg-PH1* to determine their origin. Subsequently, potential target and slice sites in the transcriptomes of both organisms were predicted. These predicted slice sites were compared to the observed slice sites in the degradome data. Based on this workflow, the sRNA analysis revealed 8,876 unique *Bd21-3* sRNAs and 12,682 unique *Fg-PH1* sRNAs. Using TAPIR about 100,000 unique intra- and cross-species mRNA targets in each of the four silencing directions were predicted (*Bd* → *Bd*, *Bd* → *Fg*, *Fg* → *Fg*, and *Fg* → *Bd*; Tab. S6, Fig. 2A−D). Interestingly, many predicted internal silencing events had a very high minimum free energy ratio (mfe-ratio) of 1, which was hardly observed for the predicted cross-kingdom interactions (Fig. 2A−D). The median mfe-ratio of the predicted interaction of *Fg*-mRNAs was slightly lower (0.63; Fig. 2B & C) than for *Bd21-3* mRNAs (0.65; Fig. 2A & D). To confirm respective mRNA slice sites in the Degradome-seq data, adaptors and restriction recognition sites of reads were trimmed, collapsed, and aligned to the transcriptomes of both organisms. After quality filtering, we aligned 154.98 million reads from 3 uninfected and 160.71 million reads from 3 infected samples to the *Bd21-3* and *Fg-PH1* transcriptomes (Tab. S2).

**Figure 2.**
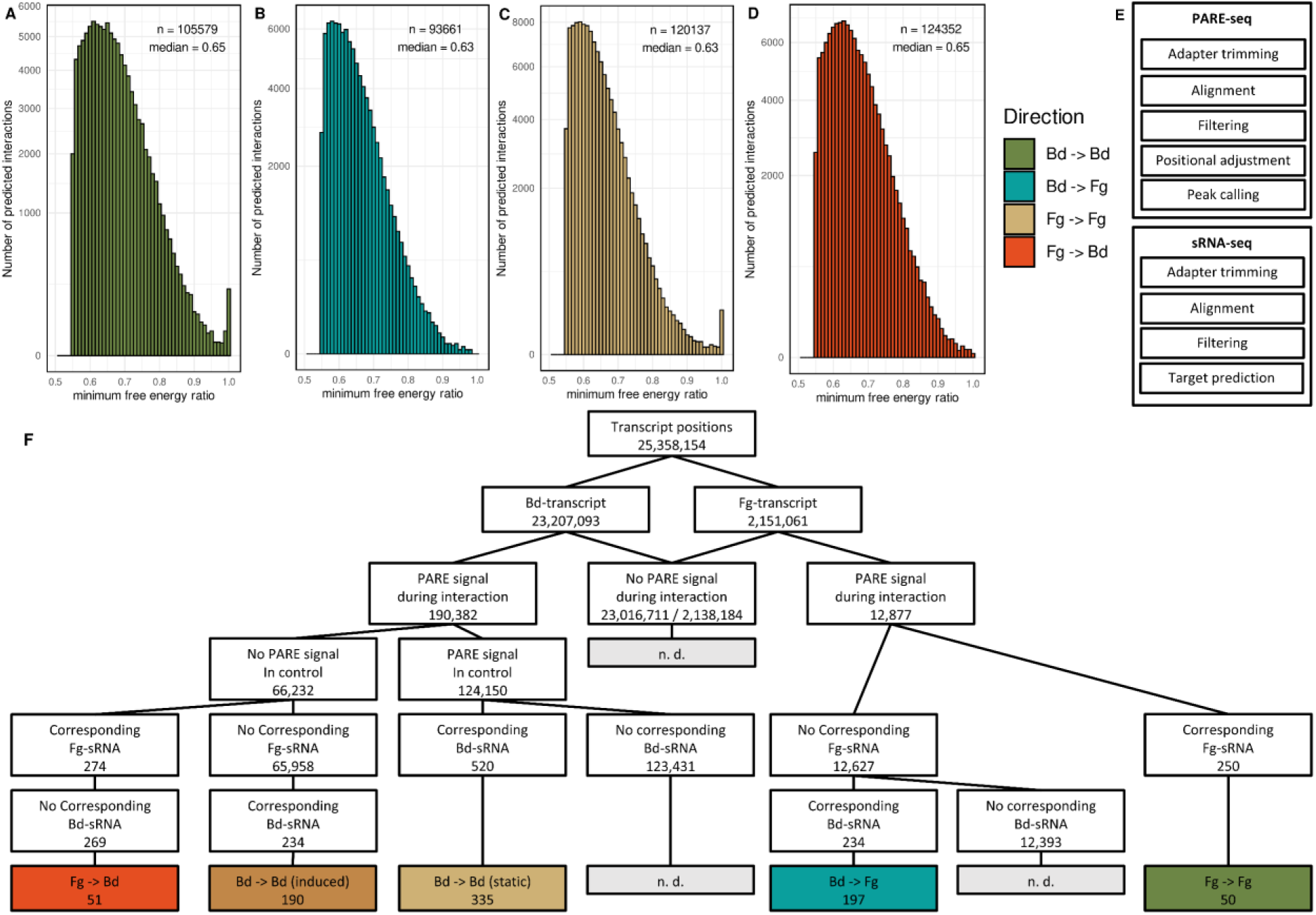
Bioinformatic identification of internal and ckRNAi-based slicing events. (A−D) The miRNA-target prediction algorithm TAPIR unambiguously assigned sRNAs, with more than 10 rpm per respective organism, to targets in the transcriptomes of both interaction partners. The histograms indicate the number of predicted unique interactions binned by mfe-ratio (binwidth = 0.01). Y-scale is square root-transformed. (E & F) Graphic representation of the ckRNAi degradome analysis pipeline. (E) Individual preparation steps and (F) decision tree of the ckRNAi analysis pipeline to identify (or not; n.d.; not defined) cleavage sites corresponding to internal and/or external sRNAs for each transcript position in the interacting organisms. Categories were defined based on the sRNA-producing and receiving organism (->). Degradome-seq data from control plants allowed discrimination between internal silencing events in *Bd21-3* that happened prior to (static) or upon infection (induced).

Perfect alignments were then filtered by strandedness, fractional counts were calculated, and reads of each transcript were normalized by z-score transformation. Since PARE techniques can create bias for reads at the 3’-end of mRNAs, a generalised additive model was calculated for all transcripts with more than 100 reads per kilobase (rpkb) and counts were corrected accordingly. Transcripts with less than 25 fractional rpkb were excluded from further analyses. Finally, slice sites were called at transcript positions with a z-score degradome intensity of at least 2 in all three replicates.

### ckRNAi analysis identifies natural *Bd21-3* sRNAs silencing *Fg-PH1* transcripts

Our study confirmed internal silencing events for 525 Bd-sRNAs in *Bd21-3* (*Bd* → *Bd*), of which 190 were exclusively found in infected samples, and internal silencing events for 50 Fg-sRNAs in *Fg-PH1* during the infection of *Bd21-3*. Even more interestingly, the analyses confirmed a total of 248 cross-kingdom (ck) sRNAs. 30 *Bd21-3* slice sites were only explainable by fungal activity of 51 sRNAs and 133 slice sites in the fungal transcripts were caused by 197 *Bd21-3* sRNAs (Fig. 2F; Tab. S7). Among the confirmed slice sites in the *Bd21-3* transcriptome were, for example, four internal target sites of miR396 in the *growth-regulating factors* (*GRFs*) *GRF5, Pto-interacting protein 1, GRF4, and GRF8* (Fig. 3A−D). It further included the infection exclusive slice site of Bdi-miR156 targeting *Bd21-3 squamosa promoter-binding-like protein* (*SPL*) *19*, a known pathogen responsive regulator (Fig. 3E) as well as one ck silencing event in *Bd21-3 lysine-tRNA ligase* by rRNA-derived *Fg-PH1* sRNA2694 (Fig. 3F). Among the confirmed slice sites in the *Fg-PH1* transcriptome were, for instance, the rRNA-derived sRNA6428 targeting *FGSG_05784*, a *Oleate hydroxylase* (Fig. 3G).

**Figure 3:**
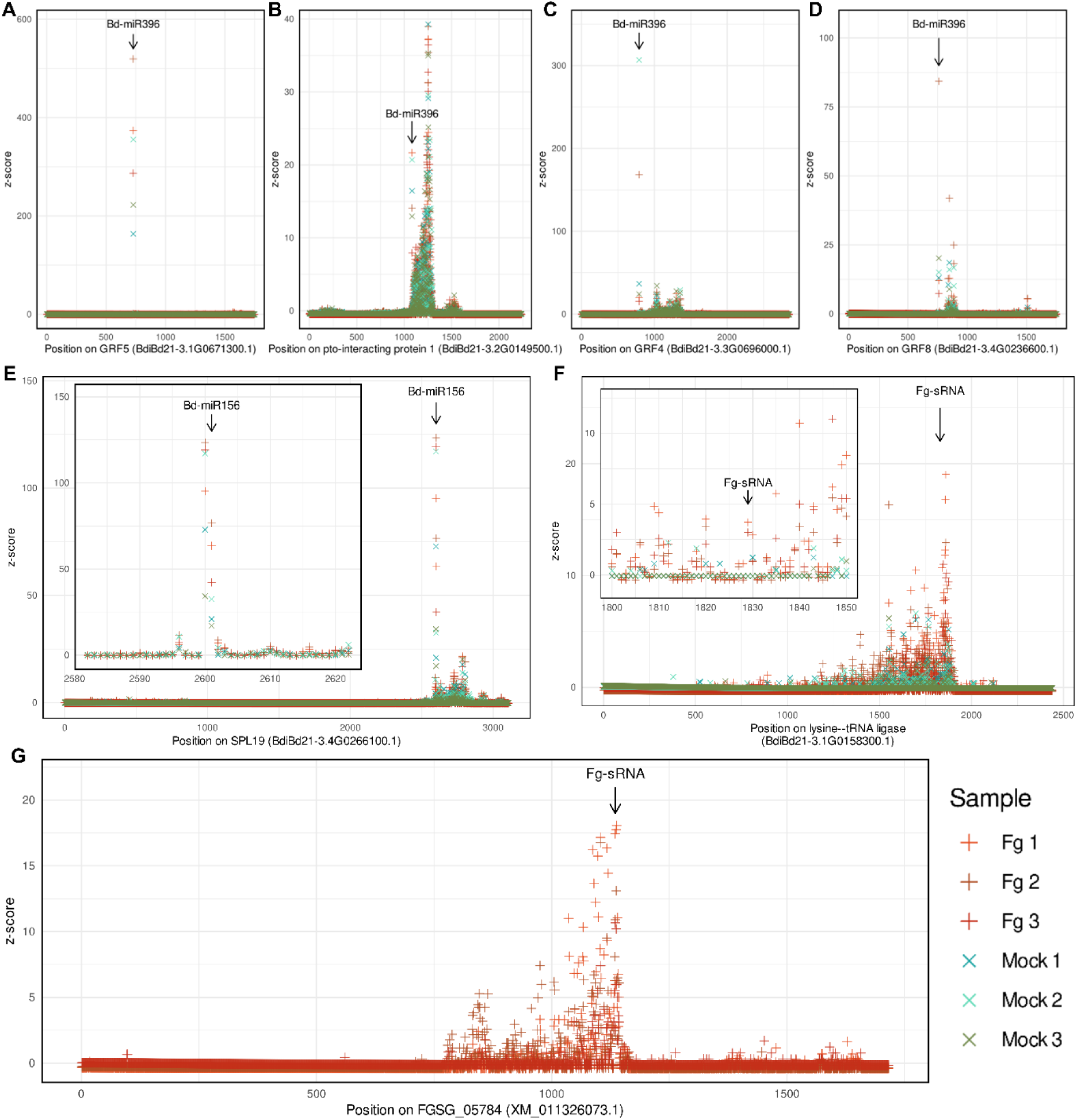
t-plots of selected silencing events which were identified with the new ckRNAi degradome analysis pipeline. Each plot shows the Degradome-seq signal strengths as z-score for each individual sequencing run for mock (x) and *Fg-PH1*-infected samples (+). Arrows indicate canonical slice site for the respective sRNAs. (E &F) Inlayed subplots enlarge specific t-plot areas for clarity

### Degradome simulations reveal sRNA pattern-to-function categories

We initially conducted a slice site analysis with CleaveLand4 (Addo-Quaye et al., 2009), a standard tool for the confirmation of slicing events from PARE data. While this approach predicted abundant active sRNAs, a subsequent simulation run with randomly generated sRNAs resulted in similar prediction outcomes. It suggests a high false discovery rate based on a random intersection of true Degradome-seq signals and predicted slice sites (Tab. S8-S16). Any random set of Degradome-seq signals and random predicted slice sites would produce a considerable overlap:

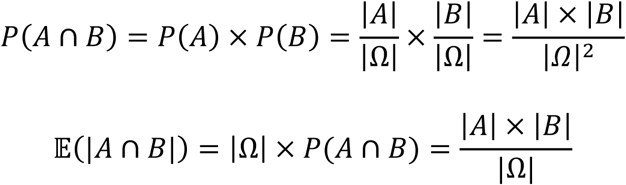

To assess a false positive rate for our data sets, a new ckRNAi degradome analysis approach was developed that considered the complex (and uneven) distribution of slice sites and target predictions and the mutually exclusive targeting of sRNAs in a cross-kingdom context. For each individual transcript, not only the true degradome data sets, but also 2000 randomly generated degradome data sets, were evaluated against the target prediction. Random degradome data sets were generated with the same number of peaks, but on random positions. The median number of confirmed slice sites in the simulations gave a sound estimate for the expectancy (E-value). The number of simulations which produced more or equal confirmations (N_s≥r_) than the real (R) data divided by the number of total simulations (N_s_) estimated the significance of respective categories.

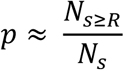

To estimate the number of true positive interactions, the median confirmations in each category (x̃_s_) were subtracted from the actual observed confirmations (R) in the respective category.

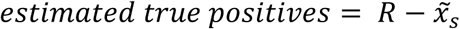

As mentioned above, the 5’-terminal nucleotide and the length of sRNAs determines their loading into Argonaute) and 21 nt 5’-U sRNAs are preferentially loaded into AGO1 in *A. thaliana* (Mi et al., 2008), explaining most currently known slice sites. This is reflected in the simulation data, where some classes outperformed all 2000 simulations, while other classes did not deviate from the expected number of confirmations in random degradomes (Tab. S17).

Overall, the simulations indicated significantly fewer confirmed interactions for random degradomes in comparison to real degradomes. This was very pronounced for 20 nt sRNAs with a 5’-U from *Bd21-3* involved in static internal silencing, where 51 confirmed interactions in the true data sets were detected in contrast to 16 median confirmations in the simulations (Fig. 4A). While the analyses uncovered 20 nt sRNAs with a 5’-G from *Bd21-3* involved in internal silencing, no differences were observed between actual and median simulated confirmed interactions (Fig. 4B). Evaluating other 5’-nucleotide options indicated that 20 nt *Bd21-3* sRNAs guiding cleavage of *Bd21-3* mRNAs carried a 5’-U, 5’-A or 5’-C while 5’-G sRNAs were inactive (Fig. 4C).

**Figure 4:**
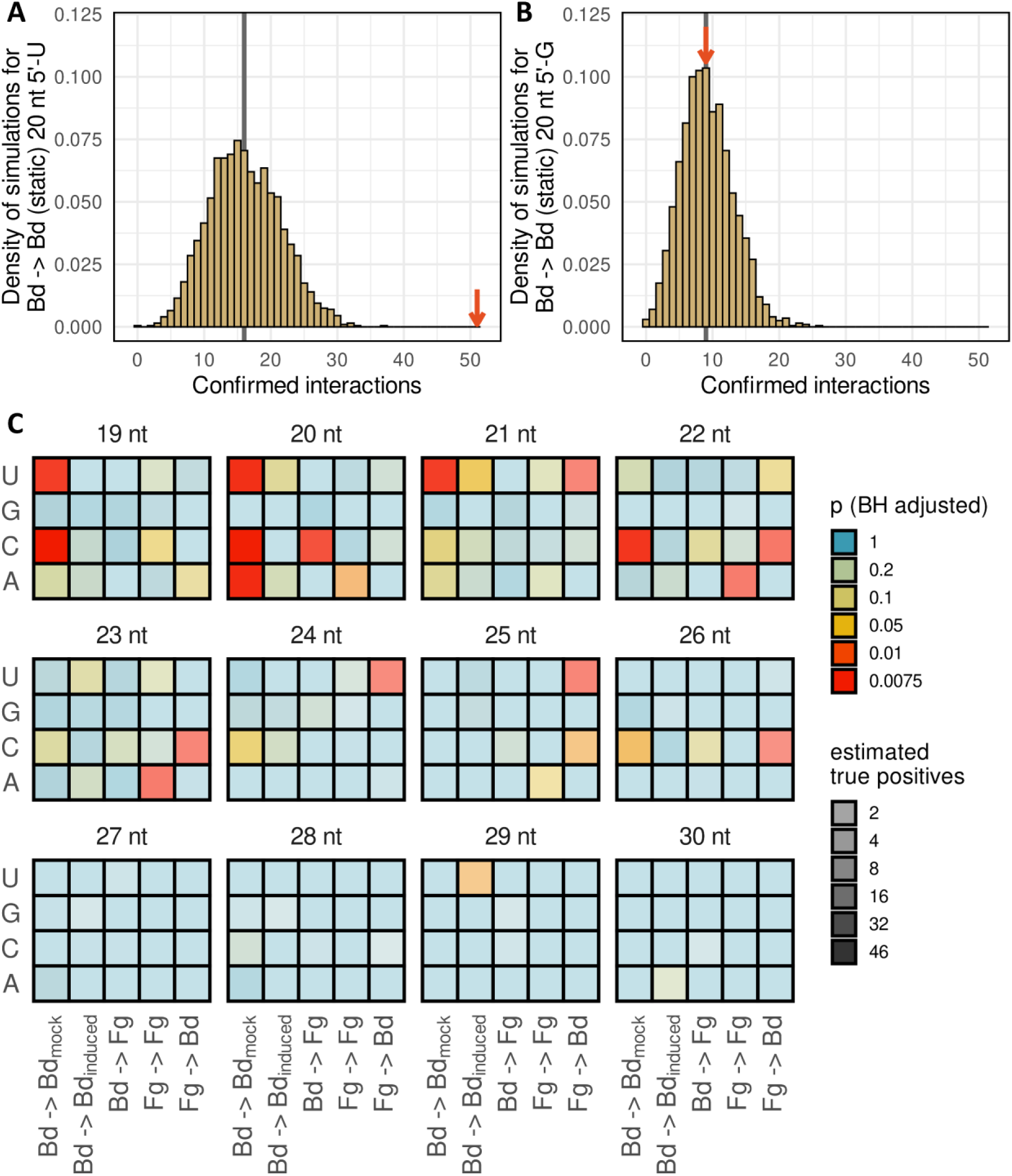
Degradome simulations indicate sRNA categories. (A-C) 2000 simulations per category and class combination evaluated the overlap of real target prediction and random degradome. (A, B) Internal (static) silencing in *Bd21-3* by 20 nt long sRNAs with either 5’-U (A) or 5’-G (B). Red arrows indicate confirmed interactions from the real degradome data. (C) Heatmaps show calculated p-values (color-coded) for all tested combinations. Red and yellow tiles show a significant enrichment of confirmed interactions in real data. Tile transparency indicates the number of estimated true positives.

Extending this simulation-based validation of cleavage to sRNAs of 19-30 nt in length and considering all four 5’-nt options revealed clear patterns. Active *Bd21-3* sRNAs with constitutive slicing activity were generally 19-22 nt long starting with 5’-A, 5’-C or 5’-U (Tab. S17; Fig. 4C). In turn, *Bd21-3* sRNA with internal activity and induced by *Fg-PH1* were mostly 21 nt long with a 5’-U. For *Fg-PH1*, sRNAs with internal slicing activity exclusively carried a 5’-A and were 20, 22 and 23 nt in length. *Bd21-3*-derived ckRNAi-sRNAs in *Fg*-PH1 and vice versa again revealed distinct patterns (Fig. 4C). Fungal ck-sRNAs silencing plant genes were 21-26 nt long with a 5’-U and/or 5’-C, while plant ck-sRNAs silencing fungal genes were 20 nt long with a 5’-C. Overall, the analyses identified 405 interactions of *Bd21-3* sRNAs with internal slicing activity (Bd -> Bd; 383 interactions under control, 22 induced in interaction with *Fg-PH1*), 22 interactions of *Fg-PH1* sRNAs with internal slicing activity (Fg -> Fg), 41 *Fg-PH1* ck-sRNAs slicing plant transcripts (Fg -> Bd), and 43 *Bd21-3* ck-sRNAs slicing fungal transcripts (Bd -> Fg) (Tab. S7 & S17). There were 219 other sRNA classes without a significant enrichment of Degradome-seq signals at the expected slice site (Tab. S17).

### *Bd21-3* cross-kingdom sRNAs mainly originate from two long non-coding RNAs and target *Fg-PH1* mRNAs regulating fundamental cellular and pathogenic processes

Our computational simulations (Fig. 4C) indicated a significant enrichment (False discovery rate (FDR) ≤ 0.0075) of slice site confirmations for active ck-sRNAs with a size of 20 nt while the group of 22 nt almost showed significant enrichment (FDR = 0.099). Both ck-sRNA groups had specific sequence characteristics. In addition to having exclusive slicing activity in the *Fg-PH1* transcriptome they are 20 or 22 nt long and carry a 5’-C. We further identified 21 nt long ck-sRNAs with these sequence characteristics. Though this group was insignificantly enriched, 21 nt ck-sRNAs were previously described to silence fungal genes (Zhang et al. 2016; Cai et al. 2018) and we therefore included them in our group of ck-sRNA candidates. In total we identified 50 unique 20-22 nt *B. distachyon* sRNAs targeting 27 different genes in *Fg-PH1* and were therefore termed anti *F. graminearum* sRNAs (αFg-sRNAs) (Tab. S18 & S19). αFg-sRNAs target *Fg-PH1* mRNAs coding for the translation machinery, such as 40S ribosomal protein S16, H/ACA ribonucleoprotein complex subunit 1, translation elongation factor 1-alpha (*EF1α*), eukaryotic translation initiation factor 1A (*eIF1A*), and small subunit ribosomal protein S35. Similarly, essential mRNAs coding for DNA replication were targeted, like cell division cycle protein 37 and DNA replication licensing factor mcm2 as well as mRNAs encoding for proteins involved in respiration (e.g. succinate dehydrogenase (ubiquinone) membrane anchor subunit). In addition to these fundamental cellular processes, αFg-sRNAs targeted fungal mRNAs participating in pathogenicity such as extracellular cell wall degrading enzymes like 42 kDa endochitinase precursor and endoglucanase 3 precursor, and components of vesicle and nuclear trafficking like AP-3 complex subunit delta, Importin-9 and a extracellular protease (alkaline protease precursor).

While each αFg-sRNA targeted only one *Fg-PH1* gene, several *Fg-PH1* genes were targeted by multiple αFg-sRNA. These αFg-sRNA were highly similar but not identical to each other and associated with the same slice site in each respective *Fg-PH1* gene suggesting an origin from the same *Bd21-3* genomic region. We therefore aligned all 50 αFg-sRNAs to the *Bd21-3* genome with the splice aware aligner STAR to gain further insights into the genomic origin and biogenesis. While some αFg-sRNAs which targeted different genes were assigned to disconnected genomic regions, all αFg-sRNAs targeting FGSG_02622, FGSG_00743, FGSG_03795 and FGSG_08811 aligned to the same genomic region (Bd3 26,537,048−26,538,548), which is highly active in the production of sRNAs (Fig. 5A & B, Tab. S19). Modelling revealed a transcript secondary structure of this locus that was untypical for a miRNA precursor (Fig. 5C), yet αFg-sRNA9 and αFg-sRNA13 were located opposite to one another while αFg-sRNA6 and 16 were separately located. Interestingly, the production of sRNAs on this locus resembled a phased pattern and the PhasiRNAnalyzer software (Fei et al., 2021) identified 21 nt phasing for this locus (Fig. 5A & B; Fig. S1). Consistent with this, two lncRNAs were annotated in this locus. BDIS_LNC000002, a 47,978 nt long lncRNA, aligned + in the same direction as 4 αFg-sRNAs (αFg-sRNA6, 9, 16, 23) while 2,578 nt long lncRNA (XR_002964634) aligned at the region in the - direction. An even larger cluster of 34 αFg-sRNAs was attributed to the BDIS_LNC005374 lncRNA which spanned several rDNA loci at the end of the chr5 rDNA repeats (Fig. S2).

**Figure 5:**
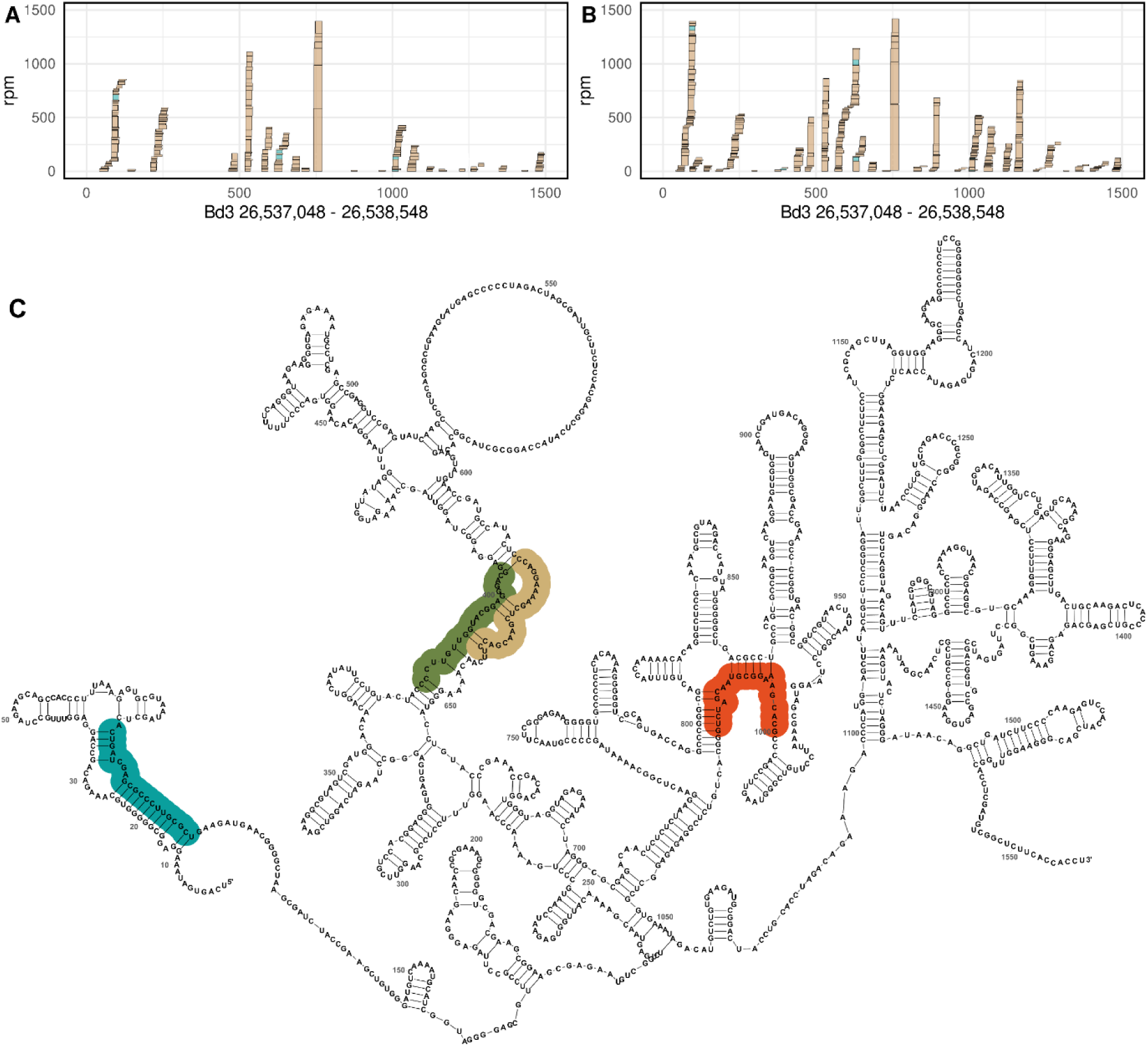
Analysis of the genomic origin of αFg-sRNAs. (A, B) Expression of sRNAs on a genomic locus in Bd21-3 control samples (A) and in infected samples (B). αFg-sRNAs are shown in blue while other sRNAs are shown in yellow. (C) Genomic locus with secondary structure producing αFg6 (blue), αFg13 (green), αFg9 (yellow), and αFg16 (red). rpm, reads per million.

### αFg-sRNA16 protects plants from *Fg*-PH1 infection in a fungal kingdom-specific manner

To assess the potential of the identified natural plant ck-sRNAs as crop protection agents, αFg-sRNA16 and αFg-sRNA23 were analysed. αFg-sRNA16, as representative of the highly slicing active 20 nt ck-sRNA group carrying a 5’C (Fig. 4C, Tab. S17), was predicted to target the fungal elongation factor 1α (*EF1α*). Though our simulation identified insignificant slicing activity for the group of 21 nt αFg-sRNA (Fig. 4C), ck slicing activity was detected for various 21 nt ck-sRNAs in different plant-pathogen systems (Zhang et al. 2016; Cai et al. 2018). We therefore included 21 nt long αFg-sRNA23 carrying a 5’C, which was predicted to target the fungal translation initiation factor 1A (*eIF1A*). To exclude unspecific effects, we designed a 20 nt long control αGFP-sRNA with a 5’-C and a similar GC content as αFg16 and αFg23 that exclusively targeted a segment of the green fluorescent protein mRNA. Considering their origin from a phased siRNA locus, all αFg-sRNA-guide strands were combined with a respective passenger sRNA with 2 nt 3’-overhangs.

For the functional assay, 2-week-old *Bd21-3* seedlings were spray-treated with sRNA duplexes of αFg-sRNA16, αFg-sRNA23 or αGFP-sRNA one day before fungal inoculation. Three days after inoculation plants were harvested and disease symptoms quantified. αFg-sRNA16 significantly reduced the relative diseased area by 37% compared to mock treatment, while αFg-sRNA23 or αGFP-sRNA showed no significant decrease in disease symptoms (Fig. 6A). In addition, foot rot symptoms were less severe in αFg-sRNA16 treated plants, and the number and severity of necrotic leaf lesions were significantly reduced (Fig. 6B). To confirm slicing activity of natural αFg-sRNA16 and αFg-sRNA23, we developed native index ligation based targeted degradome sequencing (NIL-TDS) as an independent method for Degradome-seq data. This independent analysis confirmed αFg-sRNA16- and αFg-sRNA23-mediated slicing of *Fg-PH1 EF1α* and *Fg-PH1 eIF1A* in infected but not control samples (Fig. 6C & D).

**Figure 6:**
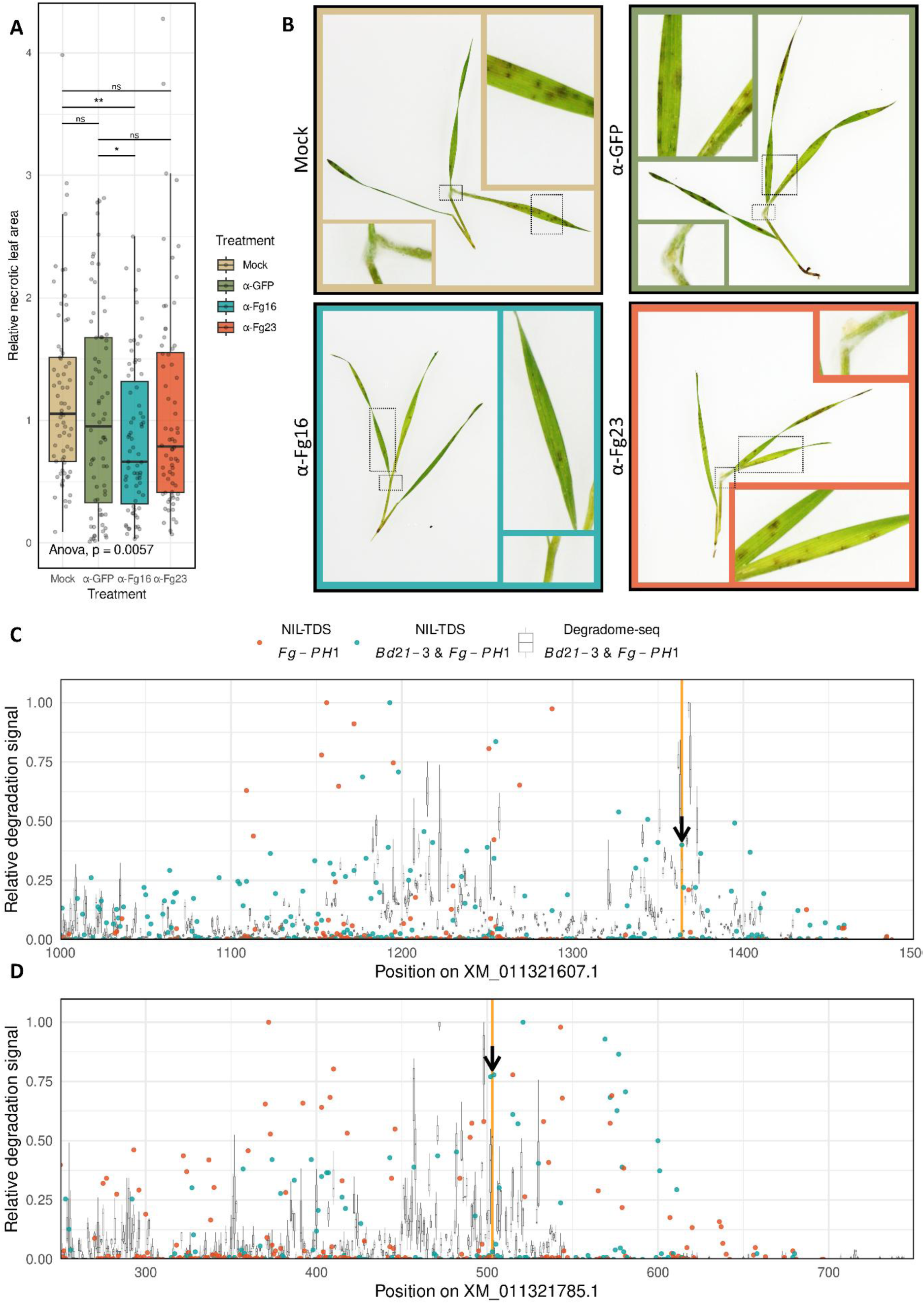
Plant protective effect of αFg-sRNAs. (A, B) *Bd21-3* seedlings treated with αFg-sRNA16 but not with αFg-sRNA23, or αGFP-sRNA show significantly reduced *Fg-PH1* disease symptoms. Measurements of mRNA degradation via NIL-TDS and Degradome-seq (C, D). Min-max normalized degradation signals from 3 infected leaf samples by Degradome-seq (gray boxplots) and infected leaf (blue) and axenically grown *Fg-PH1* (red) by NIL-TDS are shown. The expected slice sites for αFg-sRNA16 in *EF1α* (XM_011321607.1) (C) and for αFg-sRNA23 in *eIF1A* (XM_011321785.1) (D) are indicated by yellow lines.

We next wanted to know if αFg-sRNAs interact with variable or conserved sequences within target mRNAs. We hypothesized that αFg-sRNA might have undergone selection for highly conserved and, thus, invariant gene regions of pathogens during the ongoing plant-pathogen coevolution. Therefore, we aligned the target genes of all αFg-sRNAs to their respective homologs from the NCBI *Fusarium* genus reference annotations. Based on these multi sequence alignments a nucleotide conservation score was calculated. Remarkably, the target site of 74% of αFg-sRNAs was highly conserved and invariant compared to a random set of genes (Fig. 7A).

**Figure 7:**
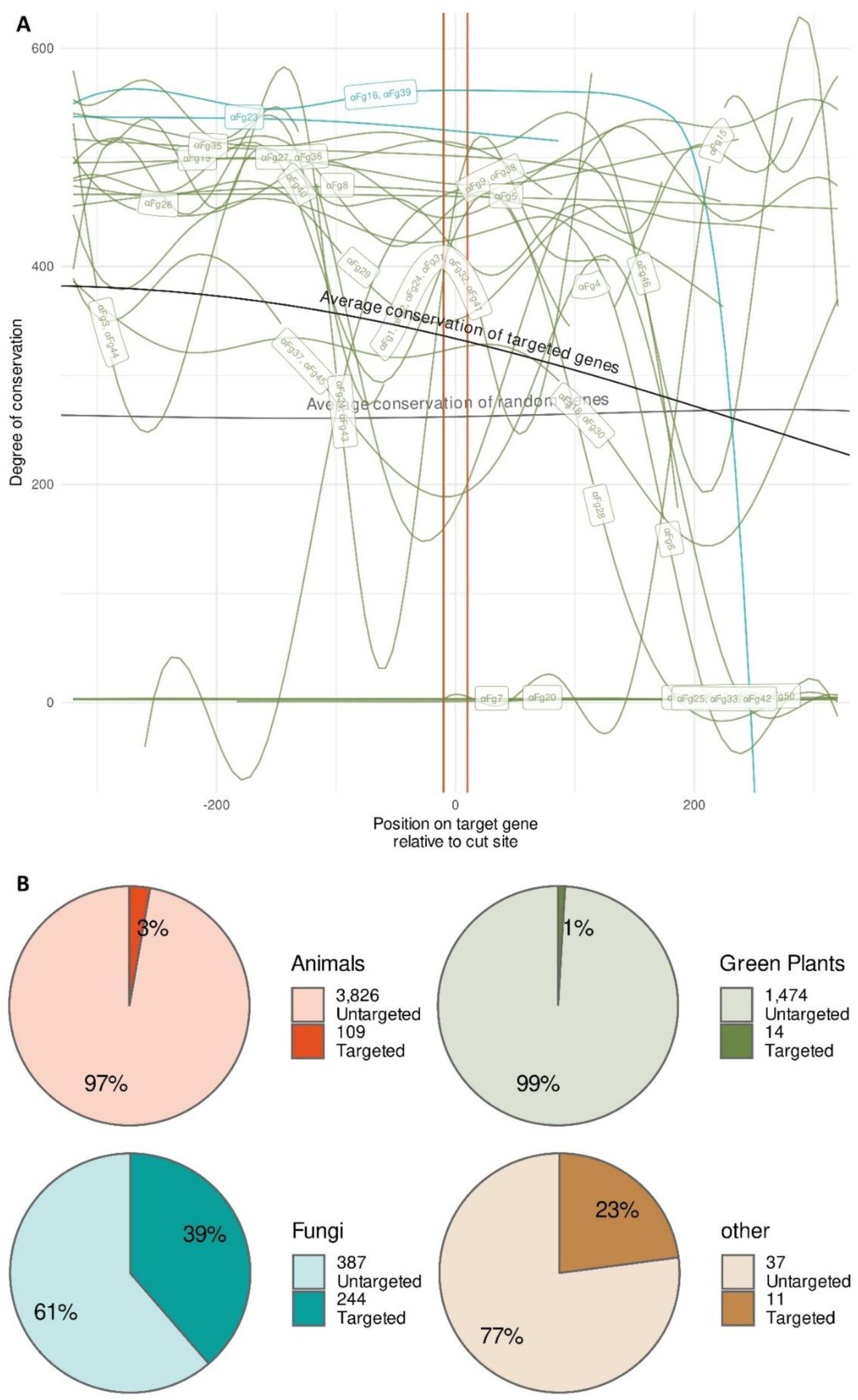
Conservation of target genes in the genus *Fusarium*. (A) The degree of conservation shows a smoothed conservation score of target gene homologs, relative to the sRNA target site (marked in red). The target genes of αFg-sRNA16 and αFg-sRNA23 are shown in blue and targets of other αFg-sRNAs are shown in green. In addition, labels indicate the targeting sRNAs for all genes. For comparison, the average conservation score of 100 random genes is shown in grey. (B) αFg-sRNA16 target site conservation in different kingdoms of life. Pie charts indicate the number of target gene homologs identified in different kingdoms. Darker colored areas in each pie indicate targeted homologs, based on computational target prediction.

Among the αFg-sRNA targets, *EF1α* showed the highest conservation among the target genes and is certainly among the most highly conserved genes in eukaryotic genomes (Fig. 7A). In fact, we found no target site variation within the species of *F. graminearum*. We therefore assessed the potential for off-target effects of αFg-sRNA16. To this end three homologs of *Fg-EF1α* from the model species *Saccharomyces cervesiae*, *Arabidopsis thaliana* and *Mus musculus* were selected. Using these representative mRNA sequences, 6,116 homologs were identified in the NCBI RNA reference sequence archive via NCBI blast. Against all homologs a target prediction was conducted. Interestingly, 39% of all fungal homologs but less than 3% and 1% of animal and plant homologs, respectively, were predicted targets of αFg-sRNA16 (Fig. 7B; Tab. S20). It indicated a high specificity and degree of target site conservation in the fungal kingdom and demonstrated a low potential of off-target effects of natural anti-fungal sRNA outside the fungal kingdom.

## Discussion

In the present study, we combined sRNA-seq with Degradome-seq for genome-wide identification of natural, plant-derived sRNAs, termed αFg-sRNA, that have antifungal activity by slicing genes in *F. graminearum*. This data set revealed their genomic organization in lncRNAs and an apparent selection for αFg-sRNA interacting with highly conserved and apparently invariant mRNA target sites. Importantly, it allowed to define the site of action (internal/static, cross-kingdom) of slicing active sRNAs based on specific sequence characteristics such as sRNA size and 5’ nucleotides. In this respect, we observed that size and 5’ properties of internally slicing sRNAs differed between *Bd21-3* (19-22 nt with 5’-A, 5’-C, and/or 5’-U) and *Fg-PH1* (20, 22, and 23 nt with a 5’-A). Interestingly, effective αFg-sRNAs from *Bd21-3* were almost exclusively 20 nt with a 5’-C, which did not mimic sRNAs of *Fg-PH1* with internal slicing activities (Fig. 4C). It suggests a mechanistic difference between *Fg-PH1* internal silencing and ck-silencing by *Bd21-3* αFg-sRNAs. In contrast, the two major fungal ck-sRNA-classes, 21 nt with 5’-U and 22 nt with 5’-C sRNAs, imitate internal slicing in *Bd21-3.* The transport mechanism for RNAs in the ckRNAi context is complex and there is evidence supporting transport via extracellular vesicles of naked RNAs and RNA bound to RNA-binding proteins (Nien et al., 2024). For Fg -> Bd ck-silencing, the transport of naked sRNAs would suffice, while αFg-sRNAs may depend on other plant-derived components to implement ck-silencing in *Fg-PH1*. Consistent with such a model, some monocot Argonaute proteins carry a signal peptide required for cellular export.

These degradome simulations enabled us to predict slicing active sRNA classes. We selected αFg-sRNA16 as representative of the 20 nt sRNA class with significant slicing enrichment and αFg-sRNA23 as member of 21 nt sRNAs with non-significant slicing enrichment. We included the latter as 21 nt sRNAs were formerly reported to have ck slicing activity. Interestingly, both sRNA showed slicing activity in our independent NIL-TDS analyses, suggesting too stringent simulation criteria, while only αFg-sRNA16 had antifungal activity in our pathoassays. αFg-sRNA23 targeted *eIF1A* which together with *eIF1* enables tRNA binding to the 40S ribosomal subunit (Passmore et al. 2007). A partial redundancy of both proteins during translation initiation was shown (Sinvani et al. 2015). This functional redundancy within the target could have compensated for *elF1A* degradation. In addition, RNA modifications in various sRNA classes can have effects on RNAi activity by either shaping immunomodulatory effects, like in the case of *N*^6^-methyladonsine (Imaeda et al. 2019), or has strong effects on sRNA stability, as is known for 2′-*O*-methylation (Li et al. 2005). It will be interesting to study in the future to what extent differences in nucleotide preferences and sRNA sizes contribute to specific RNA modifications and, thus, ck-sRNA slicing activities and efficiencies. The application of NIL-TDS might be particularly helpful here.

Plant-pathogen co-evolution follows a molecular key-lock principle driven by a multi-layered and complex interplay between plant and pathogen proteins (Jones et al., 2024). Similarly, one would hypothesize that target regions of native anti-fungal sRNAs evolved to prevent complementarity (Werner et al., 2022). In contrast, the target region of αFg-sRNA16 in the fungal *EF1α* sequence is highly conserved and apparently invariant. A highly and constitutively expressed and essential gene like *EF1α* may be restricted in changing its amino acid sequence for functionality and rely on a highly optimized codon-usage for most efficient protein synthesis. This may explain the targeting of essential and conserved genes by the identified αFg-sRNAs. Future analyses will reveal to what extent, or if at all, slight differences in the *EF1α* sequence from other fungal pathogens affect the slicing effectivity of αFg-sRNA16.

Our studies further revealed some insights with potential relevance for future application of non-coding RNAs as bioprotectants against fungal diseases. Spray-applied αFg-sRNA16 from *Bd21-3* had antifungal activities at very low dosages (1 ng/µl; about 500 ng/plant) (Fig. 6) indicating a dose response equal to chemical fungicides. In our previous studies with dsRNAs higher dosages (20 ng/µl) were applied to yield comparable reductions in *F. graminearum* symptoms in barley (Koch et al., 2019; Werner et al., 2020; Höfle et al., 2020). Similarly, Shuai et al. (2024) applied ∼136 ng/µl (10 µmol/L) when assaying 5 different designed sRNAs to reduce *F. graminearum* infection in wheat suggesting a higher efficiency of natural over designed sRNAs. It implies that current algorithms used for ds/sRNA design may not accurately predict all critical ck-interaction properties (e.g. accessibility of secondary mRNA structures and most notably length and 5’-nucleotide). In addition, dsRNAs are longer and processed into many sRNAs with approximately half of them computationally predicted to be effective (Koch et al., 2019). In turn, the conserved transcript target site of natural αFg-sRNAs (Fig. 7) might indicate an evolution of plant sRNAs for conserved transcript regions with high target site accessibility. As a result, the target affinity and thus slicing activity of natural αFg-sRNA16 might be higher. Future studies may consider these aspects and, in addition, the impact of host plants, target genes, and experimental setups on sRNA effectivities. In this respect, sRNA spray application of whole plants, as in this study, might be advantageous over detached leaf assays, which have been routinely used by us and other labs (Koch et al., 2019; Werner et al., 2020; Höfle et al., 2020; Shuai et al., 2024, Wang et al., 2016). In addition to its distance to field application, it avoids potentially conflictive wound responses from cut sites and can help to more accurately estimate effective application doses for intact plants.

## Methods

### Plant sample generation for sequencing

*B. distachyon* seeds of the cultivar Bd21-3 were placed on a wet filter paper in a 145mm petri dish and kept for 2 days in a fridge at 4°C. For germination, the petri dish was stored in darkness at room temperature (RT) for 3 days. 4 plantlets were transplanted diagonally into 10 cm square pots on F-E type soil (LD80, Fruhstorfer Erde, Germany) at 16/8 h light/dark at 22°C and a relative humidity (RH) of 60%. 14 days after transplantation, seedlings were sprayed with 10-day-old *F. graminearum* PH-1 spores, grown on SNA with a 12 h photoperiod at 25°C in an incubator, at a concentration of 500,000 spores/ml in 10 ppm Tween 20 and 0.5% gelatine. Per pot roughly 1.25 ml spore suspension was evenly sprayed on the seedlings with a compressor connected NS 19/26 thin-layer chromatography sprayer (Lenz Laborinstrumente, Germany, Cat. No. 5470004). After spraying, pots were rotated 180° and treatment was repeated. After spray application pots were placed in closable transparent plastic boxes, covered with wet tissue to keep RH close to 100%. Infection progressed in prior mentioned growth conditions with the first 24 h in darkness. 3 days post inoculation (dpi) the third most upper leaf was harvested and shock frozen. Per biological replicate and treatment 48 leaves were collected.

### Library prep and sequencing

Plant material was crushed with mortar and pestle and a phenol-chloroform extraction with TRIzol (ThermoFisher, Cat. No. 15596-026) following standard protocol was conducted. sRNA-libraries were prepared with the NEXTFLEX Small RNA-Seq Kit v4 with UDI (Perkin-Elmer / Revvity, Cat. No. NOVA-5132-32) according to manufacturer instructions. The starting material was 2 µg RNA and 16 PCR cycles used during enrichment.

For the Degradome-seq libraries the protocol of Li et al. (2019) was adapted to be compatible with NEXTFLEX UDI adapters. To this end 5’ RNA adaptor and the 5′ adaptor primer was changed to UCUUUCCCUACACGACGCUCUUCCGAUCUCAGCAG and TCTTTCCCTACACGACGCTC, respectively. This allowed the utilization of UDI adapter primer in NEXTFLEX style for the PCR enrichment step. For each library 35 µg of RNA were used.

For genome-wide analysis of gene expression, RNA sequencing libraries from isolated mRNA were generated and sequenced by the Institute for Lung Health (ILH) – Genomics and Bioinformatics – at the Justus-Liebig-University (JLU) Giessen (Germany). A total amount of 1000 ng of RNA per sample was used to enrich for polyadenylated mRNA followed by cDNA sequencing library preparation utilizing the Illumina^®^ Stranded mRNA Prep Kit (Illumina) according to the manufacturer’s instructions. Library quality was checked by capillary electrophoresis (4200 TapeStation, Agilent). Libraries were sequenced on the Illumina NovaSeq 6000 platform generating 50 bp paired-end reads.

### Bioinformatics

#### Differential gene expression

All libraries were quality controlled with fastQC (v0.12.1; Andrews et al., 2010). For differential gene expression analysis mRNA reads were filtered with cutadapt (v4.7, Python 3.12.7; Martin, 2011) to a maximum expected error of 0.5 and first two bases of R1 and first base of R2 were trimmed. Afterwards, paired reads were aligned with the splice aware aligner STAR (v2.7.11b; Dobin et al., 2013) to the Bd21-3 and Fg-PH1 reference genome assemblies (BdistachyonBd21_3_537_v1.2; Brachypodium distachyon Bd21-3 v1.2 DOE-JGI, http://phytozome.jgi.doe.gov/; GCF_000240135.3_ASM24013v3). Fragments per read were then counted with htseq-count (v2.0.5; Putri et al., 2022) and differential gene expression was analyzed with the R (v4.4.1; R Team, 2024) package DEseq2 (v1.44.0; Love et al., 2014). Differential gene expression results were plotted with ggplot2 (v3.5.1; Wickham, 2016), ggpubr (v0.6.0; Kassambara, 2023) and the palettes of wesanderson (v0.3.7; Ram & Wickham, 2023).

#### Small RNA analysis

sRNA library adapters were trimmed with cutadapt and filtered to a minimal length of 18 and a maximum expected error of 0.5 and aligned to the respective reference genomes with bowtie2 (v2.5.4; Langmead, 2012) in very sensitive end-to-end mode with penalty scores of 1,000. Alignments were filtered (concordantly aligned) and loaded into R with the GenomicAlignments package (v1.40.0; Lawrence et al.,2013) and sorted alignments were collapsed via seqtk (v1.4-r122; Shen et al., 2016) and BioSeqZip (v0.0.0.0; Urgese et al., 2020). Plotting and differential expression was conducted as described above for reads with ≥ 20rpm.

#### CleaveLand4

Degradome-seq library adapters and the anchored restriction site were trimmed and reads were filtered to a maximum expected error of 0.5, a length between 25 and 29 and “trimmed-only” with cutadapt. The trimmed Degradome-seq and collapsed sRNA reads were run in CleaveLand4 (v4.5; Addo-Quaye et al., 2009) with mode 1. To test the reliability of CleaveLand4 in a multi organism scenario, the dinucleotide frequency of all sRNA sets was calculated and random sRNAs were generated with the sample function and the package seqinr (v1.0-2; Charif & Lobry, 2007).

#### ckRNAi analysis workflow

##### Degradome peak calling

To improve degradome peak calling reads were aligned to the respective transcriptomes with STAR, allowing a maximum multimapping of 10. Aligned reads were sorted and filtered with samtools (v1.20; Danecek et al., 2021) by flag 16. Alignments were read into R with Rsamtools (v2.20.0; Morgan et al., 2021) and further filtered to again only allow strandedness + and a perfect alignment. Afterwards fractional read counts were calculated with R and pbmcapply (v1.5.1; Kuang et al., 2022). To account for positional biases in the Degradome-seq data, fractional counts were z-normalized for each transcript with more than 100 reads per kilobase (rpkb) with the outliers package (v0.15; Komsta & Komsta, 2022). To reduce computational demands, a randomly selected fifth of the data was used to model a generalized additive model of signal strength and transcript position (Fig. S3).To not be biased by slice sites, all z-scores above 3 were excluded from modelling and modelling was conducted with the “gam” function of the mgcv package (v1.9-1; Wood, 2017). The model was used to correct degradomes for all transcripts with more than 25 rpkb.

##### Target prediction

To facilitate the degradome analysis pipeline sRNA reads were treated as described above and filtered to either 2 or 10 organism specific rpm. For all sRNAs a target prediction with TAPIR (Bonnet et al., 2010) was conducted with a score cut-off of 8.5 and a minimum free energy ratio of 0.55. To enable the prediction of multiple targets of a single sRNA on the same transcript, target prediction was repeated for each gene and sRNA with the first target site removed, and results were collected accordingly.

##### Cleavage analysis

The degradome and target prediction data were combined to conduct simulation studies and confirm predicted interactions. To this end annotated RNA sequences of Bd (BdistachyonBd21_3_537_v1.2.transcript.fa) and Fg (GCF_000240135.3_ASM24013v3_rna.fna) were loaded via the seqinr package (v4.2-30; Charif & Lobry, 2007) and gene lengths for all transcripts in the degradome data were extracted. For each target gene the number of applicable peaks was calculated and saved to a vector, enabling the simulation of random degradomes. These random degradomes were analyzed with the actual target prediction data with a fuzziness of ±1, for each sRNA size and 5’nt class individually. A function of the analysis workflow was created with sRNA length as argument to allow parallelization with the parallel package (v4.4.1). The respective R script is provided as supplementary with reproducible data as RDS-file for the Fg-*EF1α* gene (Supplementary File S1, S2 & S3).

#### Biogenesis of αFg-sRNAs analysis

αFg-sRNAs were aligned to the Bd21-3 reference genome with STAR with a multimapping limit of 50. Subsequently, the genomic regions with alignments (±1kb) were written to fasta files with seqinr and all sRNA reads were aligned to these regions with bowtie2 in very-sensitive end-to-end mode with all penalties set to 1000 and a score-min of L,0,0. Alignments were plotted with a custom function in ggplot2. For this cstack size limit was adjusted to 65,536.

Phasing was analysed with the PhasiRNAnalyzer software (https://cbi.njau.edu.cn/PPSA/scripts.php; Fei et al., 2021) in a virtual conda environment at default settings. RNA secondary structure of genomic region was created via RNAcentral R2DT (v2.0; Sweeney et al., 2021) and annotated with RNAcanvas (Johnson & Simon, 2023). The genomic region of αFg16 was further used in Sequence search on RNAcentral, NCBI (Sayers et al., 2022) BLAST (Camacho et al., 2009) and the EnsemblPlants (Yates et al., 2022) BLAST to identify transcripts and further annotate the genomic region. The EnsemblPlants BLAST tool and viewer were used to visualize the retrieved transcripts via alignment.

#### Image analysis

Images from pathogenesis assays of sprayed plants and microscopic images of spores treated with sRNAs were loaded into R with the pliman package (v3.0.0; Olivoto, 2022). For plant images the background was segmented with the inverted NB index, by incrementally decreasing the cut-off value by 0.015 until the relative background size did increase by less than 0.25%. Diseased leaf area was identified via the HUE index and a threshold of 0. From each plant side 2 pictures were taken and the mean relative disease area was used for further analysis. Similarly, microscopic spore images were incrementally filtered with the inverted R index in pliman and objects were analysed via the GRAY index.

#### Target prediction for *EF1α* homologs

The annotated homologs of *EF1α* were downloaded from NCBI RefSeq (O’Leary et al., 2016) for *S. cervesiae* (NM_001178466.1), *A. thaliana* (NM_100668.3) and *M. musculus* (NM_010106.2). These transcripts were used to find homologs via NCBI BLAST. Identified homologs were downloaded with NCBI Batch Entrez and targets were predicted as described earlier.

### Spray assays

To assess the effect of αFg-sRNAs on pathogenesis the identified genomic region was utilized to generate sRNA duplexes with 2 nucleotide 3’-overhangs. These sequences were ordered at Eurofins Genomics (eurofinsgenomics.eu) as HPLC purified unmodified RNA oligos. The sequences of the αGFP-sRNAs guide and passenger strand were CGA CGU AAA CGG CCA CAA GU and UUG UGG CCG UUU ACG UCG CC respectively. RNA oligos were diluted to 100 pmol/µl in DEPC-treated water. 20 µl of each oligo, 40 µl DEPC-treated water and 20 µl 5x RNAse-free annealing buffer (300 mM KCl, 30 mM HEPES pH 7.5, 1 mM MgCl_2_) were combined in a Lobind 1.5 ml Eppendorf tube and heated for 1 min to 90°C. For proper annealing oligos were slowly cooled down over a 90 min period and subsequently kept at 4°C and were used within 24 h. *B. distachyon* plants were generated as prior described. Instead of spore solution, plants were sprayed with annealed RNA oligos at a concentration of 1 ng/µl in Gelatine [0.5 %] Tween 20 [10 ppm] solution as described earlier for spore spraying. Sprayed plants were put in plastic boxes at ambient temperature and light and lids were closed after drying. 24 h after RNA spraying, plants were inoculated with 200.000 spores/ml Fg-PH1 in gelatine Tween solution as described earlier. Plants were kept in plastic boxes at 100 %RH at ambient temperature and light and disease symptoms were documented 3 dpi.

### Native Index Ligation-based Targeted Degradome Sequencing

Native Index Ligation-based Targeted Degradome Sequencing (NIL-TDS) builds upon the method of RLM-RACE for slice site detection (Llave et al. 2002), but is followed by a mid-throughput Nanopore sequencing step. Briefly, 4µg RNA from axenically grown Fg-PH1 or from Bd21-3 infected with Fg-PH1 were mixed with 1.5 µg of RLM-adapter (5’-GCUGAUGGCGAUGAAUGAACACUGCGUUUGCUGGCUUUGAUGAAA-3’) and incubated ligation reaction was conducted with the NEB T4 RNA Ligase 1 (M0204) followed by reverse transcription with Invitrogen SuperScript II Reverse Transcriptase, and a nested PCR reaction with ThermoFisher Platinum Taq, following standard protocols using adapter-specific outer (GCTGATGGCGATGAATGAACACTG) and inner (CGCGGATCCGAACACTGCGTTTGCTGGCTTTGATG) primers and gene specifc primers ATTTGCACAAGATCCCAGGCT and ATTTCGTCGTAGAAGCGAGTACA respectively. PCR products were sequenced on the ONT Flongle platform.

## Data availability

Raw sequencing data is available at NCBI SRA via the BioProject PRJNA1307254.

## Funding

This work was funded by the Federal Ministry of Research, Technology and Space (grant BMFTR FKZ_031B1226A to P.S.) and by the Deutsche Forschungsgemeinschaft (DFG) in the research unit 5116 (exRNA) (grant number Bi 316/20-1 to PS).

## Author contributions

The work was conceptualized by P.S. & B.T.W., data was curated by B.T.W., T.P.-K. & J.W. formal analysis was conducted by B.T.W., funding was acquisited by P.S., Methodology was designed by B.T.W., S.N., R.S., T.P.-K., J.W. & P.S., Project was administered by P.S., Biological and bioinformatic resources were managed by B.T.W., S.N., T.P.-K., J.W. & P.S., Software was designed and managed by B.T.W., validation was conducted by B.T.W. and P.S., visualization was conducted by B.T.W., Manuscript was written by B.T.W. and P.S., Reviewing and editing was done by all authors.

## Supplementary Figures

**Figure S1:**
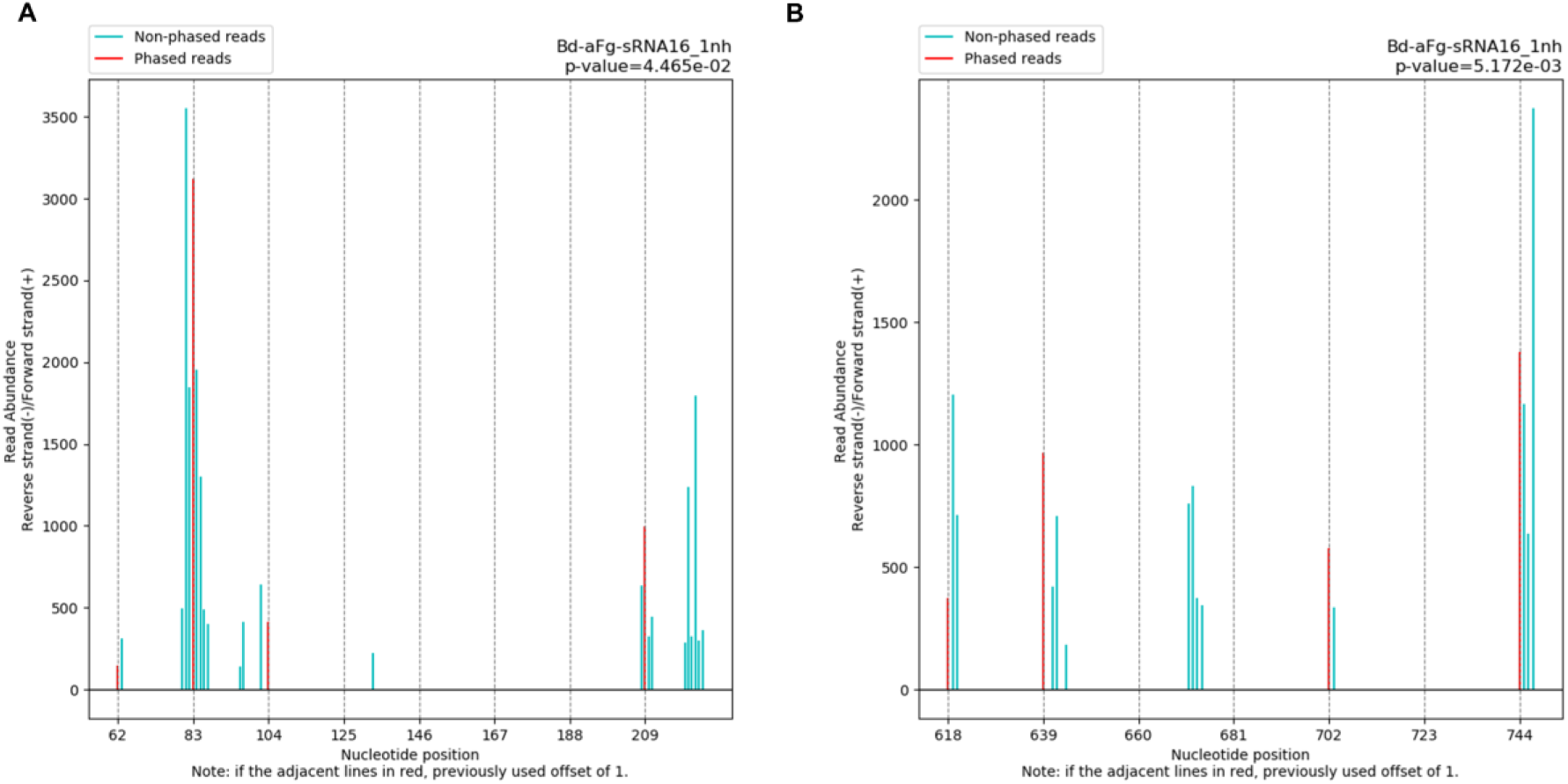
PhasiRNAnalyzer plots. sRNA alignments on genomic region around of Bd-αFg-sRNA16 show reads matching a 21-nt phasing pattern. Read start abundance of each position is shown by bars. Bars in red represent reads matching the generation of 21-nt long phased sRNA generation.

**Figure S2:**
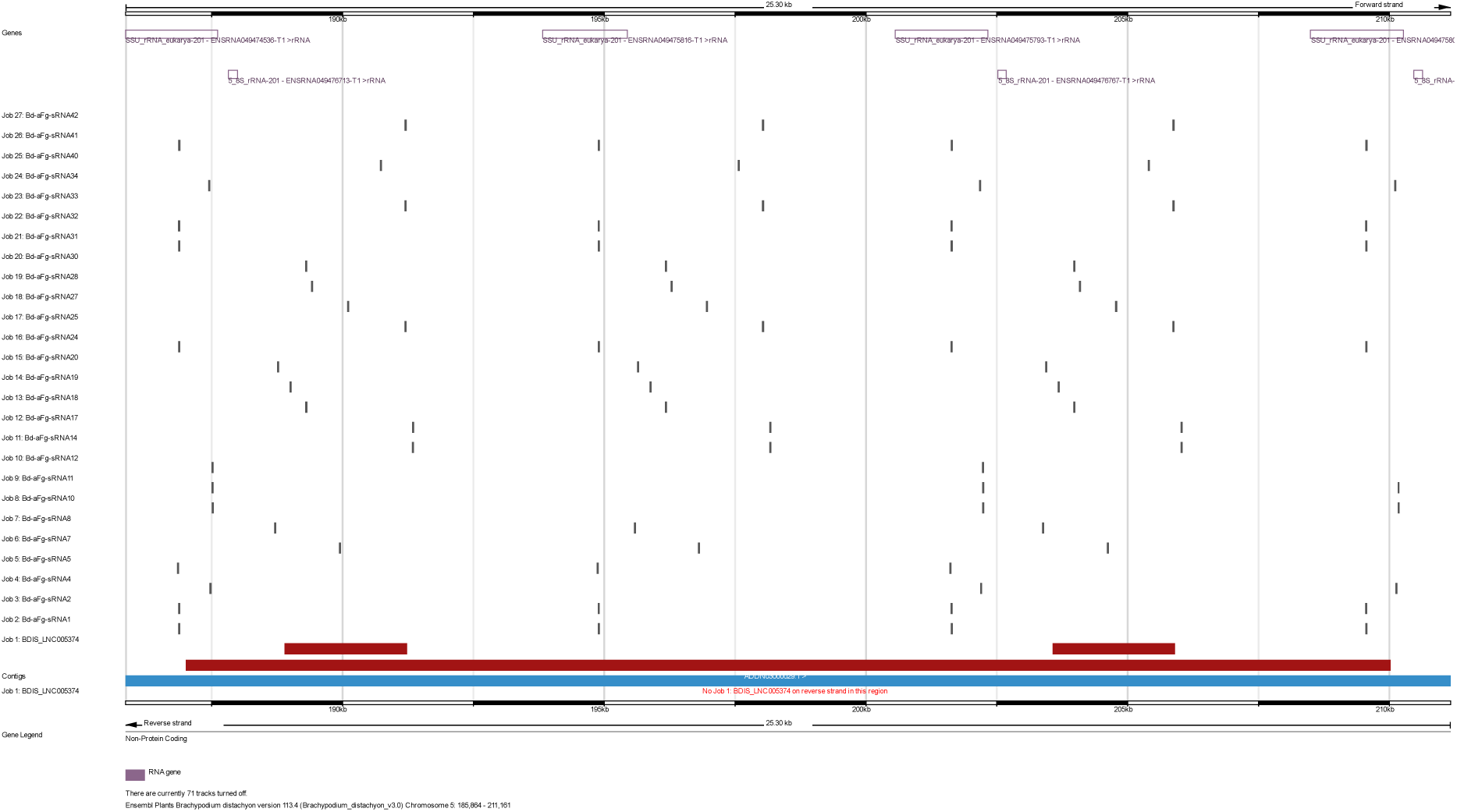
Alignment of Bd-αFg-sRNAs to the rRNA repeat locus on Bd-chr5. Here the alignments of Bd-αFg-sRNAs on a rRNA repeat locus of Bd21 is shown in the Ensembl Plants browser (Yates et al. 2022).

**Figure S3:**
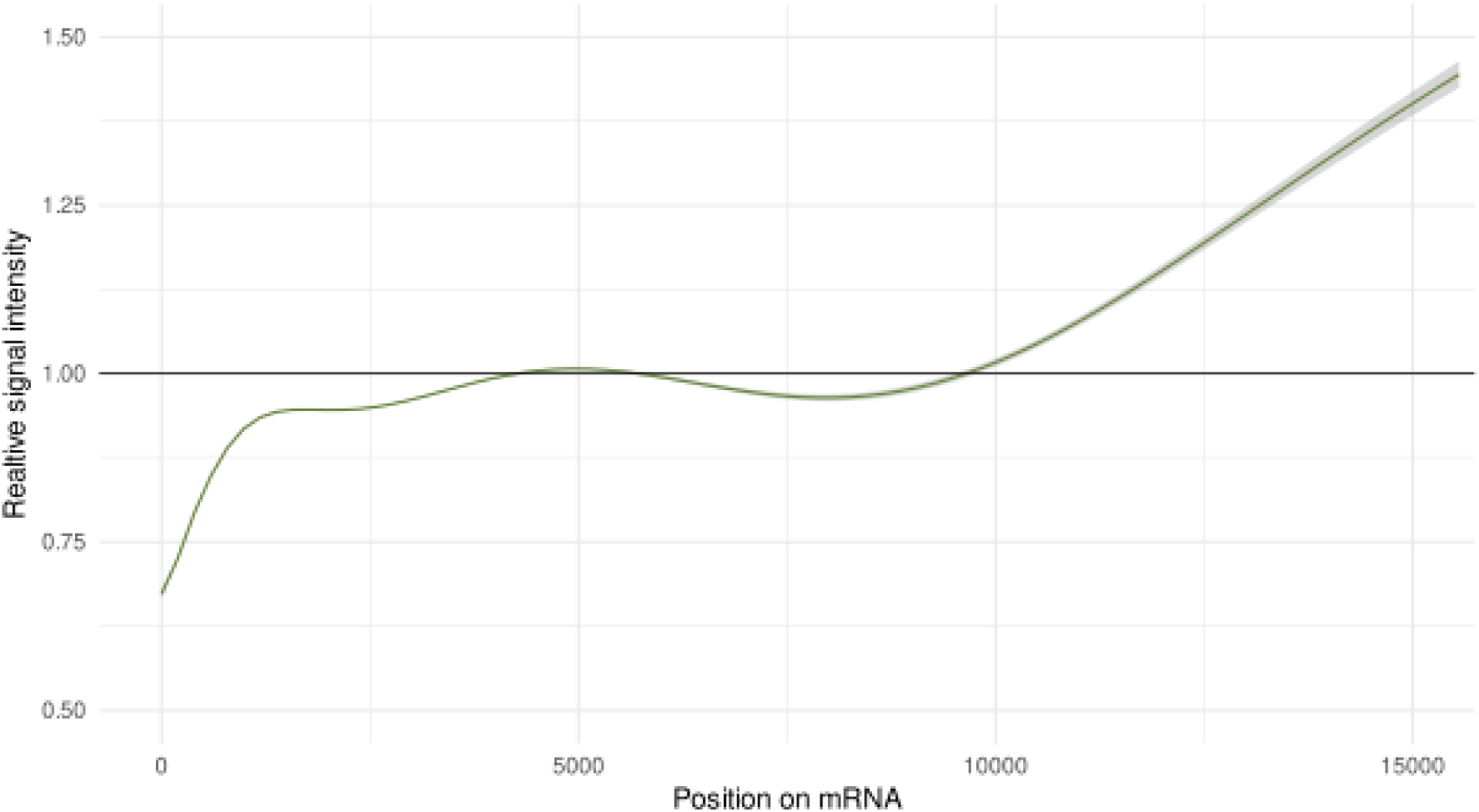
Relative degradome signal intensity. The plot relates the relative signal intensity to its respective transcript position. A randomly chosen fifth of all degradome signal intensities was used to generate this generalized additive model. The grey area around the green line indicates the 95%-confidence interval.

## Notes

### Competing Interest Statement

The authors have declared no competing interest.

